# Losing maternal care: Neotenic gene expression in the preoptic area of avian brood parasites

**DOI:** 10.1101/349118

**Authors:** Kathleen S. Lynch, Lauren A. O’Connell, Matthew I. M. Louder, Anthony Pellicano, Annmarie Gaglio, Angell Xiang, Christopher N. Balakrishnan, Eva K. Fischer

## Abstract

Parental care for is critical for offspring survival in many species. However, parental behaviors have been lost in roughly 1% of avian species known as the obligate brood parasites. To shed light on molecular and neurobiological mechanisms mediating brood parasitic behavior, brain gene expression patterns between two brood parasitic species and one closely related non-parasitic Icterid (blackbird) species were compared. Our analyses focused on gene expression changes specifically in the preoptic area (POA), a brain region known to play a critical role in maternal behavior across vertebrates. Using comparative transcriptomic approaches, we identified gene expression patterns associated with brood parasitism and evaluated two alternative explanations for the evolution of brood parasitism: reduced expression of parental-related genes in the POA versus retention of juvenile (neotenic) gene expression. While we did not find evidence for large scale gene downregulation, expression patterns did reflect substantial evidence for neotenic POA gene expression in parasitic birds. Differentially expressed genes with previously established roles in parental care were identified. Targeted examination of these selected candidate genes in additional hypothalamic regions revealed species differences in gene expression patterns is not POA-specific. Together, these results provide new insights into neurogenomics underlying maternal behavior loss in avian brood parasites.

## 1. Introduction

Parental care is an ancient social behavior that has evolved repeatedly and independently across taxa, likely because it increases offspring survival in the face of external pressures such as high predation or unpredictable access to resources (1–3). While parental care improves offspring survival it comes at a cost to the individuals providing the care (1, 4). The costs and benefits of performing parental care likely contributed to the evolution of widely varied forms of parental care within and between taxa, as well as between individuals and sexes (1, 5). Diversity in parental care strategies across the animal kingdom presents possibilities to explore the underlying genetic architecture of this behavior. While remarkable divergence in the magnitude of parental care within and between species provides fertile ground to study this behavior, animals that use an evolutionarily derived parental care strategy, rather than a strategy that is ancestral to their group, are particularly intriguing and may provide unique insight into the genetic architecture of parental care.

In birds, offspring care occurs in many forms including bi-parental care, uniparental care, and forms of cooperative breeding where siblings provide care (6–8). However, roughly 1% of the approximately 10,000 species of bird identified to date are brood parasitic, an evolutionary derived strategy in which males and females display no parental care whatsoever (9). Obligate brood parasites have completely lost parental behaviors including constructing nests, incubating eggs, and provisioning their young. Instead, they leave their eggs in the nest of another species, often with considerable negative impacts on the reproductive success of the host species. Consequently, avian brood parasites reap the benefits of parental care without any of the costs. Evolutionary losses in avian parental behavior have occurred seven independent times with three origins among cuckoos, and one origin each in cowbirds, honeyguides, Old World finches, as well as a South American duck (10–12). Brood parasitic behavior has arisen in other taxonomic groups, most notably invertebrates and fish (13–15), but it is particularly intriguing in birds as there are multiple independent events that have led five avian families across the globe to experience this substantial divergence from the ancestral state.

Behavioral ecologists have provided multiple excellent explanations for the evolution of brood parasitism including extreme limitation of nesting sites and/or dilution of the negative impacts of nest predators by not putting all one’s eggs in a single nest (9, 16–19). On the other hand, genetic and neurobiological perspectives for the appearance of avian brood parasitism are entirely lacking. The bulk of what is known about the neurobiological basis of brood parasitic behavior comes from a single published abstract which presented a study of region-specific prolactin receptor abundance (20). Prolactin receptor abundance was quantified in the preoptic area (POA), a region central to maternal behaviors across nearly all vertebrates that display parental behavior (21–24). Both the POA and the medial preoptic area (POM) are rich in steroid, peptide, and neurotransmitter receptors, all of which modulate maternal behavior (21). In brown-headed cowbirds (*Molothrus ater*), an obligate brood parasite ubiquitous across North America, prolactin binding sites within the POA exhibit reduced sensitivity (20) as compared to redwinged blackbirds (*Agelaius phoeniceus*), a closely related blackbird that is not brood parasitic (Fig. 1A). On the contrary, prolactin itself is not significantly reduced in brown-headed cowbirds as compared to red-winged blackbirds (25), indicating the difference between these related species with stark divergence in parental care is in region-specific prolactin receptor expression but not circulating prolactin. As prolactin permits parental care in several vertebrates, including both male and female birds (26–30), this initial investigation suggested that brood parasites may serve as natural knockdown system in which to understand maternal care and the loss of this behavior in 1% of avian species.

**Figure 1.**
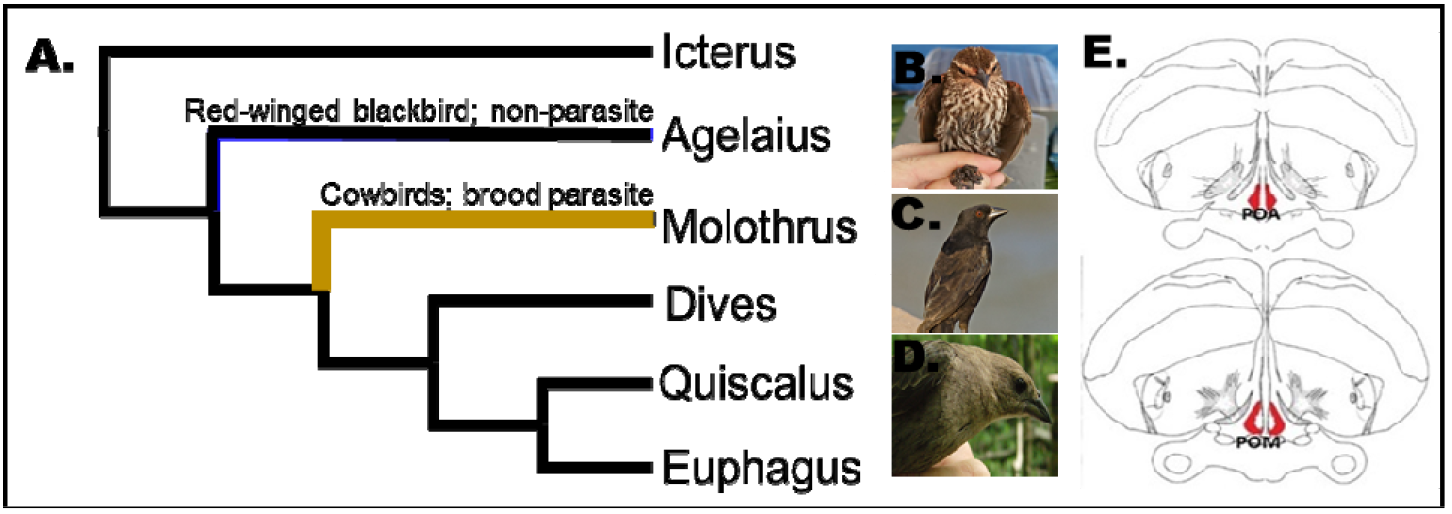
Overview of the Icterid genera (i.e. blackbirds) and target brain regions. (A) Blackbird phylogeny including the three focal species: (B) red-winged blackbird (*Agelaius phoeniceus*), (C) bronzed cowbird (*Molothrus aeneus*) and (D) brown-headed cowbird (*Molothrus ater).* The brown branch of this phylogeny represents the only brood parasitic genus within the Icteridae family. (E) Illustration of the brain regions used for transcriptome assembly and gene expression analysis. The red regions represent the brain regions collected. POA = preoptic area; POM = medial preoptic area.

The evolution of brood parasitic behavior represents, at least in part, a failure to transition from a non-maternal to a maternal state. Here, we explore this failure to transition into a maternal state by exploring the genomic basis for brood parasitism using brain region-specific gene expression comparisons. These comparisons address two non-mutually exclusive explanations: natural “knockdown” in brood parasites of genes associated with maternal care versus retention of juvenile (neotenic) gene expression in the POA. Because brood parasitism is a derived behavior that likely evolved from a maternally-caring ancestor, brood parasites presumably once possessed the necessary molecular and neurophysiological machinery to produce an operational “maternal brain”. Thus, it is possible that whole suites of genes, rather than just prolactin receptors, have been “knocked down” through evolution in brood parasites. Alternatively, brood parasites may retain neotenic gene expression patterns in the POA. Neoteny is a developmental pattern that occurs when adults retain juvenile characteristics, and therefore does not require novel mechanisms or genetic variants, but rather relies on the alteration of existing mechanisms via shifts in developmental timing. Neoteny in plumage color and skull ossification patterns has been described for brood parasitic birds (31), so a compelling hypothesis is that whole suite of neotenic traits are underpinned by broad-scale neoteny in gene expression. Novel morphologies and functionalities commonly result from modifications in timing of gene expression during development (32–34), and here we examine whether loss of maternal care in brood parasites is regulated by a shift toward neotenic gene expression.

In the present study, we examine alternative mechanisms for the evolution of brood parasitism by taking advantage of divergence in maternal care within North American Icterids (i.e. blackbirds) to identify POA-specific gene expression differences associated with the evolution of obligate brood parasitism (Fig. 1B). We compare two brood parasitic Icterids, brown-headed and bronzed cowbirds (*M. aenus*), with a non-parasitic Icterid, the red-winged blackbird. Our aims are three-fold: (1) identify POA-specific changes in gene expression associated with brood parasitic behavior, (2) explore whether differentially expressed genes follow a pattern of knockdown or neotenic gene expression, (3) identify candidate genes for future studies exploring mechanisms of brood parasitic behavior.

## 2. Materials and Methods

### (a) Bird capture and treatments

Female brown-headed (N = 14) and bronzed cowbirds (N = 17) as well as adult female red-winged blackbirds (N = 4) and female juvenile red-winged blackbirds (N = 5) were captured in Texas using mist nets and bait traps in May-June of 2014 and 2015. All female bronzed cowbirds and red-winged blackbirds laid eggs in their cage the morning after capture. The brown-headed cowbird females displayed adult plumage but no eggs were noted in these cages. All birds were transported to Hofstra University in a climate-controlled space with three birds per cage (16.5" L x 11.8" W x 22" H). Birds were then housed in outdoor aviaries for two weeks. During transport and housing, birds were fed a modified Bronx Zoo diet with mealworm supplements. All procedures listed here were reviewed and permitted by Hofstra University IACUC (#13/14-18).

Cowbirds exhibit elevated prolactin levels during the breeding season even though they do not exhibit parental care (25). Therefore, females received osmotic minipumps (Azlet, model 1007D, DURECT Corp. Cupertino, Ca.) containing prolactin while other females received saline/bicarbonate vehicle alone. Prolactin implants contained ovine prolactin (Sigma; St. Louis, Mo.; 3.3μg/hr; 80 μg/day in .87% NaCl/0.01M NaHCO3, 3:1 ratio v/v: 12μl/day) because it successfully elevated prolactin levels effect in doves (27, 35, 36). Subcutaneous implants of osmotic pumps released 0.5μl of prolactin / hour for 7 days. Pumps were implanted subcutaneously using isoflurane anesthesia. Sample sizes for each species/treatment are as follows: brown-headed cowbirds: N = 9 prolactin, N = 5 vehicle; bronzed cowbirds: N = 11 prolactin, N = 6 vehicle; red-winged: N = 4 prolactin. On day 8, females were rapidly decapitated. Brains were flash frozen on dry ice followed by −80°C storage until sectioning. Blood was collected via the trunk and immediately centrifuged for 10min at 30,000g followed by storage at −80°C until assayed. The largest follicle was measured for all subjects after sacrifice. The average largest follicle size was 0.5 mm ± 0.39 in bronzed cowbird, 1.5 mm ± 0.62 in brown-headed cowbird, 0.6 mm ± 0.40 in adult red-winged blackbirds and unmeasurable in juvenile red-winged blackbirds.

The efficacy of the prolactin osmotic pumps was verified using prolactin EIA assay kits from Aviva systems biology (San Diego, CA; chicken: 0KEEH00637) with plasma taken from large sub-set of subjects (prolactin treatment total N = 19; bronzed N = 12; brown-headed N = 7; vehicle total N = 7; bronzed N =3; brown-headed N = 4) to assess the efficacy of prolactin osmotic pumps. Prolactin samples were measured on two plates with an intra-assay variation of 8.23% and an inter-assay variation of 5.8%. The detection limit was 0.3 ng/ml and the manufacturer reported no detectable cross-reactivity with other relevant proteins. A student t-test with unequal variances was used to compare circulating prolactin between treated and untreated females.

### (b) Sectioning and sequencing

Brain tissue was sectioned into 200μm sections on a Leica CM1950 cryostat and thaw mounted onto microscope slides. Coronal sections from the caudal extent were sectioned. As the POA was approached, tissue sections were set upon a frozen, RNAse treated slide and examined under a dissecting scope containing dry ice on the stage to keep the section frozen. If the boundaries of the POA were confirmed (Fig. 1B), a 1.22 mm diameter tissue punch (Myneurolab, Leica, Richmond IL) was collected and placed into 500μl of Trizol (Fischer Sci., Waltham, Ma.). POA and POM sections from both hemispheres were collapsed into one tube and will hereafter be referred to as POA. All POA sections along the midline to the third ventricle from the caudal split in the tractoseptomesenphalicus to the rostral anterior commissure were collected. All tissue was stored at −80°C until preparation for sequencing.

RNA was isolated following Trizol manufacturer specifications and was immediately followed by mRNA isolation using NEXTflex PolyA Beads according to manufacturer specifications. RNA sequencing library prep was completed using NEXTflex Rapid Directional RNA-seq based kits. Briefly, first strand synthesis was completed via RNA fragmentation immediately followed by NEXTflex rapid reverse transcriptase. The assembled product was placed in a NEXTflex directional second strand synthesis and immediately cleaned up using Agencourt AMPure XP beads. NEXTflex adenylation mix was used for end repair on second strand synthesis DNA. Adaptor ligation was completed using NEXT flex ligation mix and NEXTflex RNAseq Barcode Adaptors immediately followed by another cleanup with AMPure XP beads. Cleaned up DNA was combined with NEXT flex Uracil DNA Glycoslylase, NEXTflex PCR Master Mix and NEXTflex Primer Mix and amplified using standard PCR. PCR product was immediately cleaned up using AMPure XP beads. POA transcriptomes were sequenced at Harvard University using an Illumina HiSeq 2500 following procedures described in (38, 39).

### (c) Transcriptome construction & annotation

We constructed *de novo* transcriptomes for each species separately following the same procedure. We used R corrector to amend Illumina sequencing errors (40). We next trimmed reads using Trim Galore! (Babraham Bioinformatics, Babraham Institute) to remove Illumina adapters and restrict all reads to only high-quality sequence. Following developer recommendations, we used a quality score of 33, a stringency of 5, and a minimum read length of 36 bp. We pooled corrected, trimmed reads from all individuals by species prior to transcriptome construction. We used Trinity (41, 42) to construct reference transcriptomes for each species. Our initial assemblies contained 296,578 contigs for brown-headed cowbirds, 394,219 contigs for bronzed cowbirds, and 243,189 contigs for red-winged blackbirds.

We filtered our raw transcriptome assemblies using several approaches. We first used used CD-HIT-EST (Weizhong Li Lab) to cluster overlapping contigs and removed any contigs that were smaller than 250bp following clustering. To remove contaminant sequences, we annotated sequences using blastx queries against the SwissProt database and retained only those contigs with annotations to known vertebrate genes. We used default parameters for our blastx queries with an e-value cutoff of 10^−10^. We chose this approach to minimize contaminants and because our primary focus in this study was on genes known to be important in social behavior (and more specifically maternal care) across vertebrates. We annotated our final assemblies using Trinotate (43) and used BUSCO (44) to assess assembly completeness based on conserved ortholog content across highly conserved vertebrate genes. This provided assembly completeness estimates at 64% in brown-headed cowbirds, 78% in bronze cowbirds, and 50% in red-winged blackbirds. These estimates are in line with expectations for single brain region transcriptomes (45). All high-powered computing for transcriptome assembly and filtering was performed on the Odyssey computer cluster supported by the FAS Science Division Research Computing Group at Harvard University.

### (d) Identification of orthologs

To compare expression across species in an unbiased way, we generated a matched set of orthologs found in each of the three transcriptome assemblies. We focused on alignment of open-reading frames to identify protein-coding genes and to ensure high-quality sequence alignment. Briefly, for each species-specific final transcriptome we predicted the ORFs for each contig using TransDecoder (42). The resulting ORF sequences were then clustered and reduced using CD-HIT (%ID = 99.5). We then used the longest ORF for a given ‘Trinity gene’ and identified orthologs present among all three species with a reciprocal best hit blast approach using blastp with an e-value cutoff of 1e^−20^ (46). The predicted polypeptides for orthologs were then aligned with MAFFT (47) and the corresponding coding sequences were back-aligned with pal2nal (48). Finally, each alignment was subjected to alignment scoring and masking using ALISCORE (default parameters) and poorly aligned regions were trimmed using ALICUT (49). This procedure left us with 12,237 aligned orthologs shared across all species. Using this list, we filtered our complete assemblies to obtain species specific transcriptomes that contained only transcripts that were matched and aligned across the three species.

### (e) Read quantification and differential expression analysis

To quantify gene expression in a manner comparable across species, we aligned reads to the species-specific ortholog assemblies. We mapped reads and estimated their abundance with Kallisto (50) using default parameters. We combined read counts into a single matrix for all individuals from all species based on ortholog identification and used this count matrix for all downstream analyses.

We then examined differential expression (DE) of orthologs using DESeq2 (51) to run standard, pairwise DE analysis for our comparisons of interest. We corrected p-values for multiple hypothesis testing using standard FDR correction and set our adjusted p-value cutoff to <0.05. We identified contigs differentially expressed (DE) in the following four comparisons: (1) between prolactin treated parasites compared to vehicle treated parasites, (2) each brood parasite compared to adult non-parasite (i.e. bronzed cowbird vs red-winged and brown-head cowbird vs redwing), (3) between brood parasite species (i.e. bronzed vs brown-headed cowbirds) and, (4) between adult and juvenile non-parasites. Following DE analysis, we performed GO term enrichment analysis for DE genes using the ‘Biological Processes’ GO categories in the topGO package (52) in R.

Behavioral transformations that occur when a female becomes a mother are associated with structural and functional plasticity throughout the brain (53). Maternally-related plasticity in hypothalamic regions is dependent on neuromodulators. However, in mammals, neurochemical and structural plasticity are dependent on one another such that critical changes in neuromodulators often induce structural plasticity in the brain of the new mother and vice versa (54–58). Therefore, we performed targeted analysis of candidate genes identified as playing a role in maternal and social behavior and categorized these genes as belonging to neuromodulators that regulate changes in existing cell function or structural genes that underlie large scale neural renovations. For this list of candidate genes, we report results from DE analysis with a less stringent p-value cutoff of < 0.1 (after FDR correction) as our study includes wild-caught birds, extremely small tissue punches, and a smaller sample size of parental species; all of which contribute to variation that may obscure biologically relevant results.

### (f) Quantitative PCR

To determine if the gene expression patterns observed in the POA were specific to this region, a subset of subjects had additional tissue punches removed during cryosectioning (N = 6 red-winged blackbirds; N = 13 bronzed cowbirds). These tissue samples were punched from four additional hypothalamic regions demonstrated to possess labelling for phosphorylated signal transducer and activator of transcription 5 (pSTAT5) in non-oscine brids (27). pSTAT5 detection is an indication of prolactin receptor activity. Additional hypothalamic brain regions with pSTAT5 labeling includes: ventral medial hypothalamus (VMH), lateral hypothalamus (LH), posterior medial hypothalamus (PMH) and the tuberal nucleus (TU). These additional hypothalamic regions were collapsed into a single sample. RNA isolation and cDNA synthesis were conducted as described in Lynch et al (59). Briefly, tissue punches were homogenized and RNA extracted using Trizol (ThermoFischer, Waltham, MA) following manufacturers instructions. Extracted RNA was DNase treated using turbo DNA-free kits (ThermoFischer, Waltham, MA). Reverse transcription of cDNA was done using Superscript First-Strand Synthesis (Invitrogen, Carlsbad, Ca.). The qPCR primers were designed using Primer3 and purchased from Sigma (St. Louis, MO). The qPCR was conducted using SYBR green detection on an Applied Biosystems StepOnePlus v. 2.2.3 (Applied Biosystems, Foster City, Ca.) with each sample run in triplicate. Gene expression levels were normalized by dividing by cDNA starting input quantities as measured by a RiboGreen RNA quantification assay (59, 60–62). We used the Ribogreen assay (Quant-iT RiboGreen RNA reagent (ThermoFischer, Waltham, MA) for accurate quantification of starting cDNA concentrations, which eliminated housekeeping gene comparisons. RiboGreen reagent measures single-stranded RNA and cDNA with equal effectiveness (61). Because cDNA quantification better reflects the actual target input into each well in the qPCR procedure, we chose to use cDNA estimates in previous studies (59) as well as this study. The estimated quantity from the qPCR results were averaged across the three replicates. We conducted individual t-tests with correction for unequal variance to compare quantification of four candidate genes selected from the transcriptome assembly. These genes include mesotocin, arginine vasotocin (AVT), galanin and prostaglandin synthase. Benjamini-Hochberg procedures were used as the FDR correction. All sequences for these genes were derived from the transcriptome assembly described above.

## 3. Results

### (a) Prolactin treatment of brood parasites

Prolactin assay kits were validated by diluting a sample with unknown prolactin concentrations into three concentrations. The slope of the line from the dilution test was compared to the slope of the line of the standard curve to assess parallelism between the slopes as described in Lynch and Wilczysnki (37). The slope of the line for the serially diluted prolactin samples was −9.6, and the slope of the line for the prolactin standard curve provided by the manufacturer was −11.

Osmotic pumps were effective at elevating circulating prolactin levels in treated females compared to blank implanted females of both species (t_19_ = 2.48; p = 0.02.). However, only three transcripts differed between treated and untreated bronzed cowbirds and only a single transcript differed between treated and untreated brown-headed cowbirds. Thus, very few transcriptome differences occurred due to prolactin treatments. Given the lack of differences in POA gene expression following prolactin treatment, we combined treated and untreated birds in all further analyses.

### (b) Gene expression differences associated with brood-parasitism

We received 498 million 100-bp paired-end reads that passed the HiSeq quality filter, averaging 12 million reads per sample. We filtered the species-specific transcriptome to generate assemblies containing only the 12,237 transcripts orthologous across all species (brown-headed N = 14; bronzed N = 17; red-winged blackbirds N = 4). To identify transcripts whose expression patterns were associated with parasitic versus non-parasitic behavior, we conducted pairwise differential gene expression (DE) analysis among our three focal species. There were 119 differentially expressed transcripts between brown-headed cowbirds and redwinged blackbirds, 634 differentially expressed transcripts between bronzed cowbirds and redwinged blackbirds, and 112 differentially expressed transcripts between bronzed and brown-headed cowbirds (Fig. 2; see S2 for complete list of DE genes). Of the transcripts differentially expressed between parasites and red-winged blackbirds, 82 were overlapping between bronzed and brown-headed cowbirds. Of these shared differentially epxressed transcripts, 81 showed expression changes in the same direction in both parasites as compared to red-winged blackbirds. We refer to these transcripts as concordantly differentially expressed (CDE; Fig. 2;listed in table S2). Shared changes in CDE genes in both parasitic cowbirds species as compared to non-parasitic red-winged blackbirds, suggest expression differences are most likely associated with the evolution of parasitic behavior.

**Figure 2.**
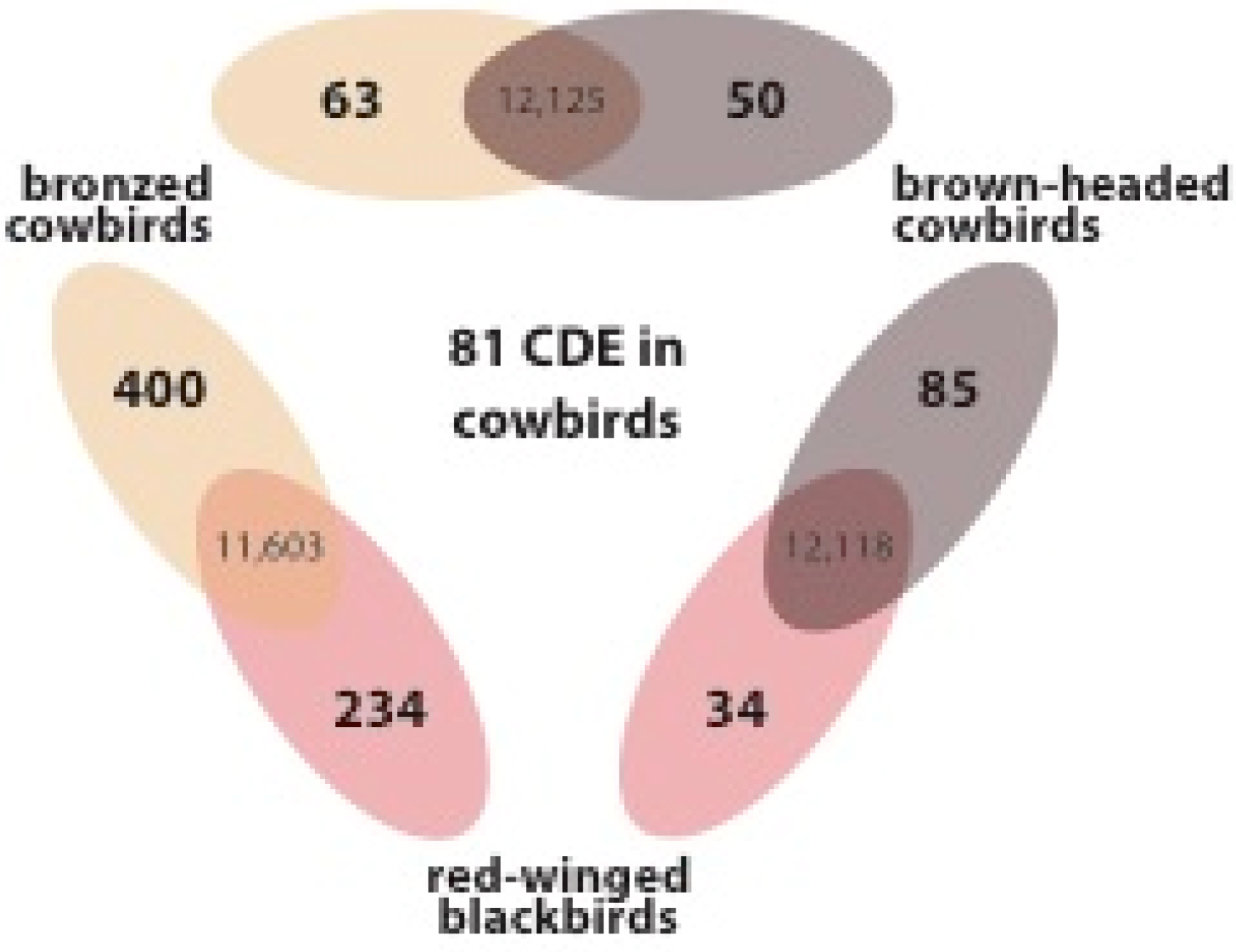
Differential gene expression comparisons between parasitic cowbirds and non-parasitic red-winged blackbirds. Transcripts differentially expressed in pairwise comparisons between species are shown in Venn diagrams with the number of differentially expressed transcripts on the edges and the number of non-differentially expressed transcripts in the overlap. The number of genes closer to a given species indicates those transcripts that were significantly upregulated in that species. 81 transcripts were concordantly differentially expressed in parasitic cowbirds, meaning they were significantly differentially expressed in the same direction in both parasitic species as compared to non-parasitic red-winged blackbird adults, suggesting these genes are those most likely to be involved in the evolution of brood parasitism.

We performed GO-term enrichment analysis for all comparisons and for the list of CDE genes (Table S3). Notably, GO categories associated with neuropeptide signaling (p = 0.008) were associated with all comparisons among parasites and non-parasites, but not with comparisons between parasites (S3).

### (c) Brood-parasitism: knockdown vs neoteny?

Two general patterns of gene expression change in the POA may be associated with brood parasitism. First, there may be downregulation (i.e. ‘natural knockdown’) in key POA genes in brood parasite. Alternatively, the parasite POA may not transition to adult-like patterns of gene expression. Instead, parasites may retain juvenile-like (neotenic) expression patterns. To address these alternative explanations we (1) examined the direction of divergence in differentially expressed transcripts between cowbirds and red-winged blackbirds, and (2) asked whether expression changes between cowbirds and red-winged blackbird adults reflected those between juvenile and adult red-winged blackbirds.

Overall, gene expression differences between parasites and red-winged blackbirds were biased toward increased expression in cowbirds relative to blackbirds: in brown-headed cowbirds 71% (85/119) of differentially expressed transcripts increased expression relative to red-winged-blackbirds, and in bronzed cowbirds 63% (400/634) of differentially expressed transcripts increased expression relative to red-winged-blackbirds. Of the CDE transcripts, 78% (63/81) increased expression in both cowbirds relative to red-winged blackbirds (Fig 2).

To address the idea of juvenile-like gene expression associated with a loss of maternal care, we additionally sampled red-winged juvenile blackbirds (N=5) and estimated transcript abundance to map to the red-winged blackbird ortholog assembly. We first compared POA gene-expression in red-winged blackbird juveniles and adults. We found only 16 genes that were significantly differentially expressed between red-winged blackbird juveniles and adults (Table S4). While there were few significant expression differences, we were nonetheless able to use the direction of expression differences in juveniles versus adults to examine whether expression differences in cowbirds were more juvenile-like (neotenic) or adult-like. To do so, we compared the direction of gene expression changes (i.e. up or down regulation) in parasitic cowbirds and juvenile red-winged blackbirds as compared to adult red-winged blackbirds. When differences between parasites and juveniles were in the same direction (i.e. both up-regulated or down-regulated as compared to adult non-parasites) we considered transcripts as exhibiting juvenile-like expression (Fig. 3A). Conversely, when differences between parasites and juveniles were in opposite directions we considered transcripts as exhibiting adult-like expression (Fig. 3A). We performed this comparison only for CDE transcripts, as these represent the set of genes most likely involved in the transition to parasitic behavior.

**Figure 3.**
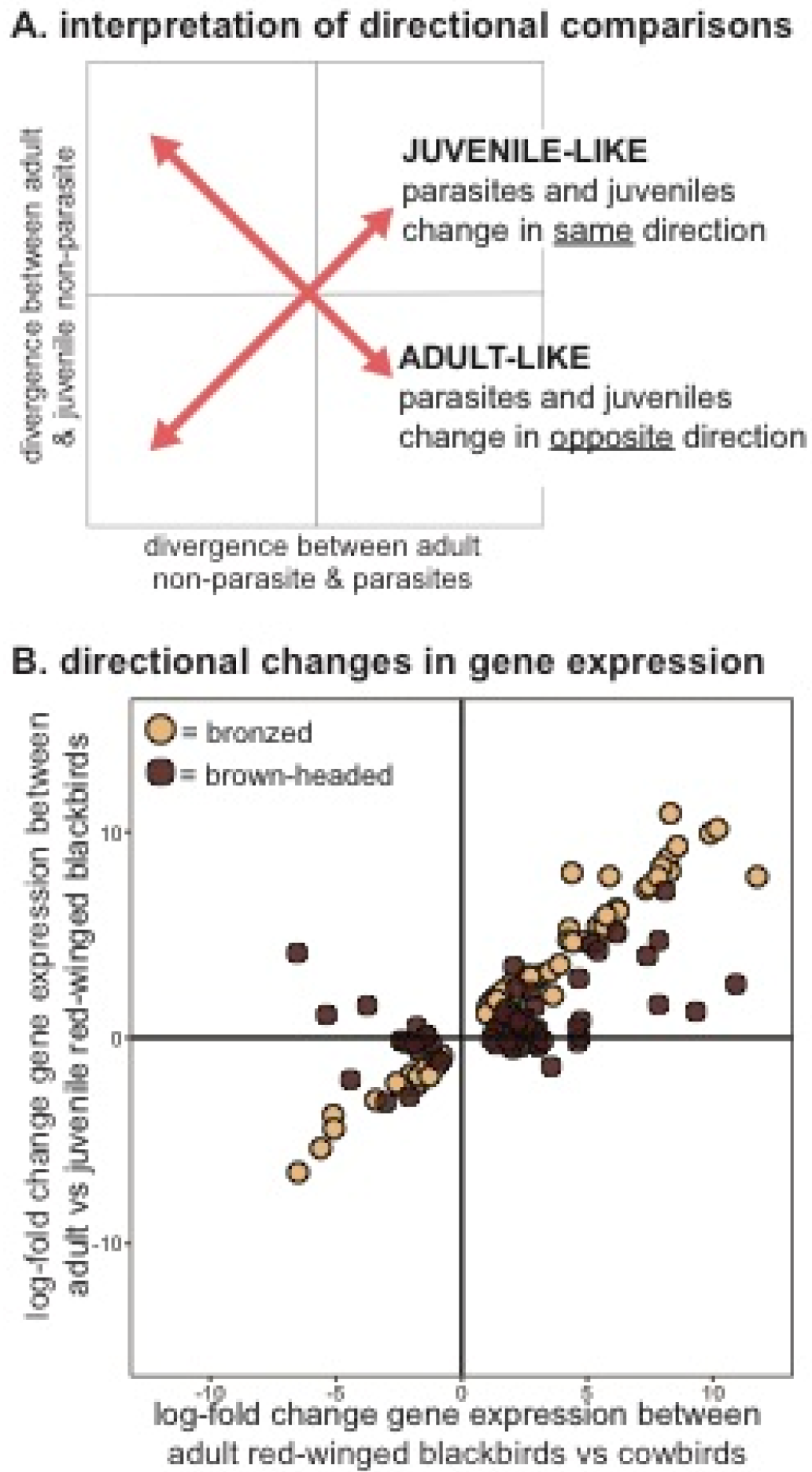
Neotenic patterns of gene expression in the POA of parasitic cowbirds. (A) Whether gene expression changes in adult parasitic cowbirds are juvenile-like (neotenic) or adult-like as compared to non-parasitic blackbirds is reflected in the directional relationship between expression differences. When parasites mirror juveniles the relationship is positive (bottom left and top right quadrant) and when parasites mirror adults the relationship is negative (top left and bottom right quadrant). (B) We find that for both bronzed (tan circles) and brown-headed (brown circles) cowbirds expression patterns are overwhelmingly juvenile like, with 78% percent of transcripts exhibiting juvenile-like expression. Data is shown for transcripts CDE in both cowbird species (i.e. those most likely to be involved in behavioral transitions to parasitism).

Of those transcripts DE between bronzed cowbirds and adult red-winged blackbird, 74% showed juvenile-like expression in the parasitic species. Similarly, 76% of genes DE between brown-headed cowbirds and adult red-winged blackbirds showed juvenile-like expression in the parasite. Finally, for the transcripts CDE among bronzed and brown-headed cowbirds, 78% showed juvenile-like expression in the cowbirds (Fig. 3B). In brief, expression differences between parasitic cowbirds and adult red-winged blackbirds largely exhibited juvenile-like expression patterns in parasitic species.

### (d) Identification of candidate genes

The induction of maternal care in new mothers is associated with structural and functional plasticity throughout the brain (53). Therefore, we used our lists of DE genes to identify candidate genes previously demonstrated as involved in both types of plasticity. We identified 38 genes with established roles in functional and structural plasticity. Twelve genes with demonstrable roles in social and parental behavior appear in figure 4. Supplemental table 5 lists all candidate genes and citations that demonstrate their role in regulation of social behavior, including maternal care. These genes include modulators that regulate changes in existing cell function including metabolism and structural genes that underlie large scale neural renovations. We included metabolic-related genes as this differential expression may be a consequence of the brood parasite never switching from a self-focused to an offspring-focused individual. Of the 38 genes identified as regulating structural and functional plasticity, 21 transcripts are neuromodulators and their receptors, 10 transcripts are involved in structural and synaptogenic modification, and 7 transcripts are peptides and receptors involved in metabolic regulation. Among these 38 candidate genes, 24 exhibit neotenic expression patterns and of these genes, 18 are CDE with juvenile red-winged blackbirds.

**Figure 4.**
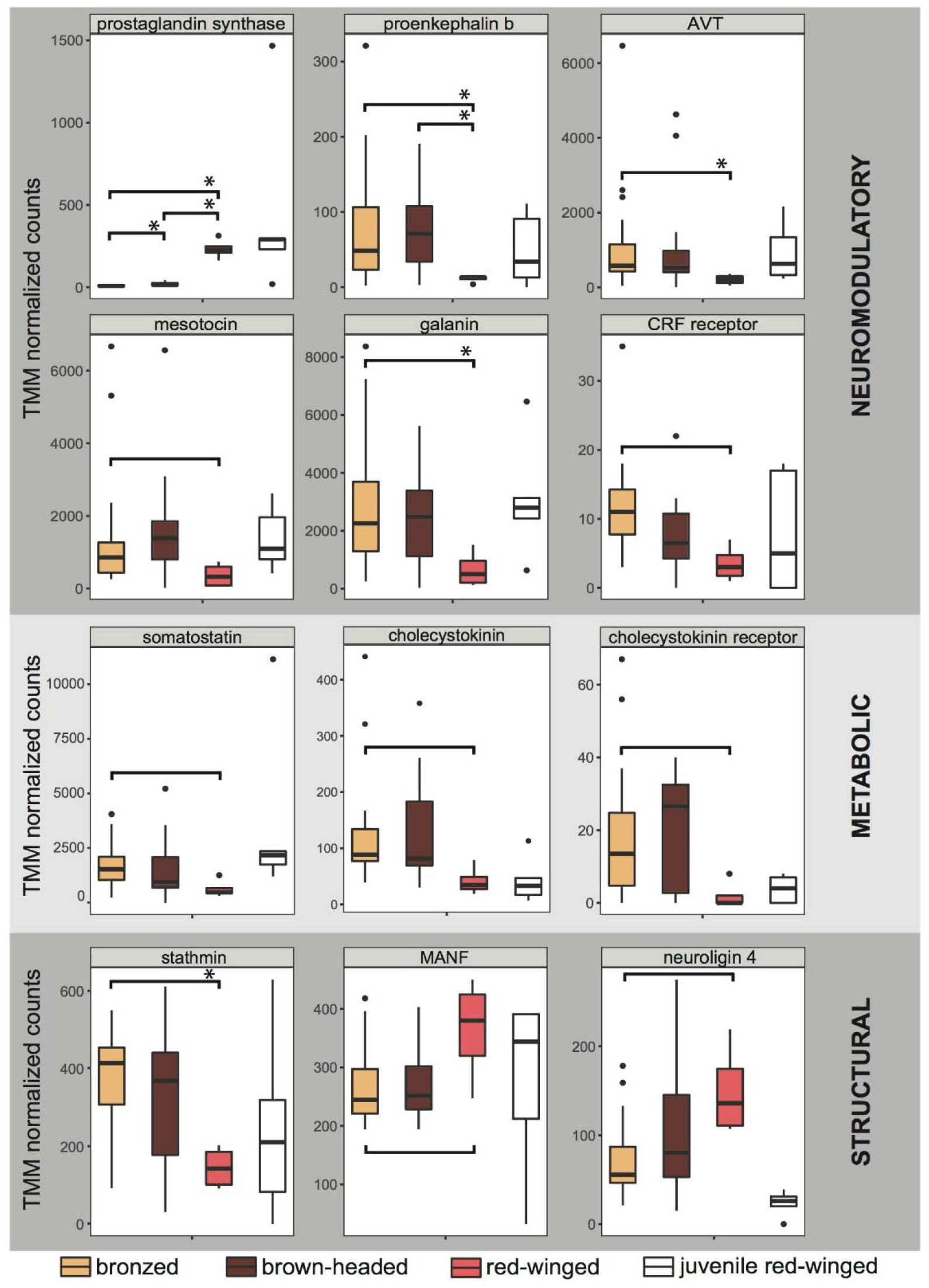
Transcript expression of candidate genes. Functional plasticity genes included neuromodulators and metabolic modulators. Structural genes included genes involved in changes to neural architecture. Select candidate genes known to be important in maternal and social behavior are presented here. Boxplots show TMM normalized transcripts counts for presumptive candidate genes in bronzed cowbirds (tan), brown-headed cowbirds (brown), adult red-blackbirds (pink), and juvenile red-winged blackbirds (white). Genes are grouped based on their known neuromodulatory (top), metabolic (middle), or structural (bottom) functions. Significant differences are indicated by brackets either with (p>0.05) or without (p>0.1) asterisks. Remaining candidate genes associated with structural and functional plasticity are listed in Table S5. AVT = arginine vasotocin; CRF receptor = corticotropin releasing factor receptor; MANF = mesencephalic astrocyte-derived neurotrophic factor

### (e) Candidate genes in other key hypothalamic regions

Four candidate genes were selected for examination in additional key hypothalamic brain regions including the ventral medial hypothalamus (VMH), lateral hypothalamus (LH), posterior medial hypothalamus (PMH) and the tuberal nucleus (TU). These regions were pooled into a single sample to determine whether differential gene expression of these genes is specific to the POA. Results reveal mesotocin and galanin are significantly elevated in bronzed cowbird hypothalamic regions relative to red-winged blackbirds (Fig. 5; t_12_= 4.81; p = 0.01; t_13_ = 2.36, p = 0. 03 respectively). Prostaglandin synthase and AVT were elevated in red-winged blackbirds compared to bronzed cowbirds but this difference was only significant for AVT (Fig. 5; t_5_= −2.43; p = 0.059; t_5_ = −3.6, p = 0.05 respectively).

**Figure 5.**
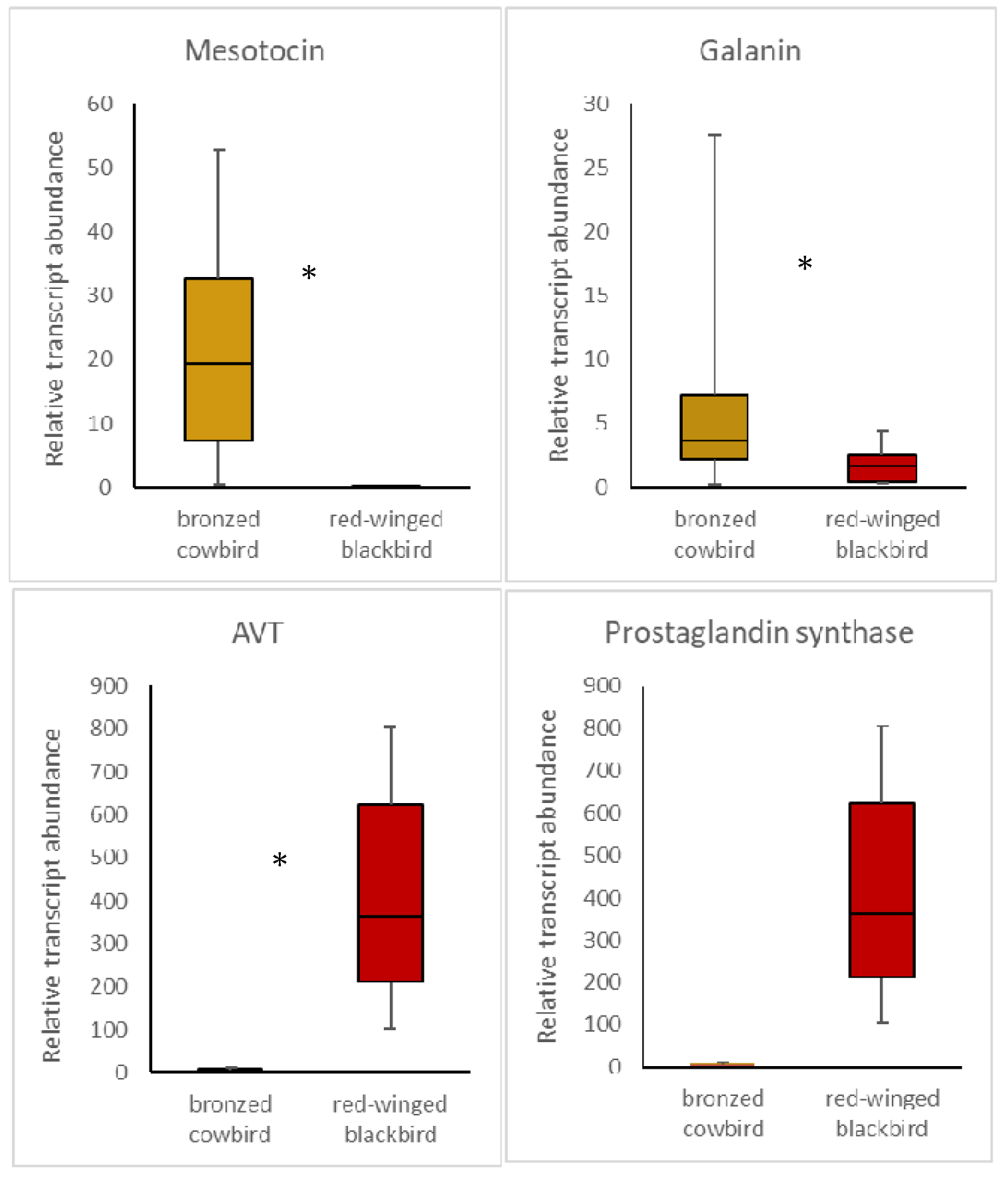
Candidate gene expression in additional brain regions. Four additional hypothalamic brain regions were punched from brain tissue and collapsed into a single sample to examine candidate genes in these regions. These hypothalamic brain regions include ventral medial hypothalamus (VMH), lateral hypothalamus (LH), posterior medial hypothalamus (PMH) and the tuberal nucleus (TU).

## Discussion

The evolution of avian brood parasitism provides fertile ground for the investigation of mechanisms mediating parental care and its loss within a powerful, comparative framework. Here, we provide the first insights into the transcriptional mechanisms associated with the evolutionary transition from maternal care to brood parasitism in Icterids (blackbirds). While behavioral ecologists have provided excellent explanations for the evolution of brood parasitism, this study is the first to address the neurobiological and molecular basis for the loss of maternal behaviors in avian brood parasites.

While there are many reasons that genes may be differentially expressed among species, the close phylogenetic relationship, similar ecology, and highly conserved function of the POA implicate those differentially expressed genes with similar expression patterns across brood parasites as those most likely related to the loss of parental care. We therefore identified transcripts that showed significant expression changes in the same direction in both parasitic cowbirds species as compared to non-parasitic red-winged blackbirds (i.e. transcripts that were concordantly differentially expressed; CDE). We identified 81 CDE transcripts, which we suggest represents the set of transcripts whose expression changes in the POA are the most likely contributors to brood parasitic behavior, and therefore provide excellent candidates for future work in avian and other vertebrate systems. Indeed, the evolution of brood parasitism in the cowbirds present in the Americas represents only one of several independent evolutions of brood parasitism in birds. Broader phylogenetic comparisons across genera will determine whether CDE transcripts identified here are implicated in convergent losses of parental care across Passeriformes. Investigating this set of genes across these genera within Passeriformes in future will provide insight into mechanisms of convergent behavioral evolution.

Following identification of the set of transcripts most likely involved in the transition from maternal care to brood parasitic behavior, we tested two alternative and non-mutually exclusive hypotheses concerning general mechanisms associated with brood parasitism. First, given the known importance for POA activation in the induction of parental behaviors across taxa (21, 63–67) and previous work demonstrating prolactin receptor down-regulation linked to brood parasitism (20), we reasoned that generalized gene downregulation might inhibit the transition to maternal behavior in brood parasites. In this case, gene expression changes should reflect a downregulated pattern (i.e. ‘natural knockdown’), especially in genes associated with the induction of maternal care. Alternatively, we hypothesized that an absence of the ability to transition from a non-maternal to a maternal state might be reflected in the retention of juvenile-like (neotenic) expression patterns in the POA. We emphasize that these patterns are not mutually exclusive as downregulation in the expression of key genes could be a component of neotenic expression patterns.

The pattern of differential gene expression is in a direction that opposes predicted patterns that would support the “natural knockdown hypothesis”. In fact, POA-specific expression patterns of DE genes were significantly upregulated in 78% of CDE transcripts in cowbirds as compared to red-winged blackbirds. We emphasize, however, that a lack of evidence for downregulation at a global level does not contradict the potential importance of downregulation of individual genes, such as the prolactin receptor or of brain region-specific downregulation of vitally important genes. Thus, further investigation is needed to understand whether gene knockdown still plays an important role in the appearance of brood parasitic strategies. An additional possible explanation for the observed pattern of gene expression is that transcription and translation are mutually independent processes that have different timings, locations, and functional complexes (68). Therefore, post-transcriptional regulation may cause a decrease in POA protein levels that is not reflected in mRNA expression. While changes in protein abundance cannot be definitively predicted from mRNA abundances in this study, our identification of CDE transcripts provides strong candidates for future functional studies that can explicitly investigate the correspondence of mRNA and protein and the behavioral consequences of expression changes across these levels of biological organization.

While overall gene-expression differences between cowbirds and red-winged blackbirds were not consistent with large-scale expression downregulation in parasites, we found marked evidence for a shift toward juvenile-like (neotenic) expression in the POA of adult parasites: 78% of CDE transcripts exhibited juvenile-like expression in parasitic cowbirds, suggesting that the retention of a neotenic, non-maternal expression state in the POA may be partly responsible for non-maternal parasitic behavior. Innovative morphologies and functionalities are known to result from modifications of gene expression timing during development, and a retention of neotenic expression patterns is known to underlie the evolution of complex traits across species (69, 70), including behavioral traits in primates (34). For example, advanced cognitive abilities of human primates are a behavioral phenotype thought to be derived from neotenic processes. Somel et al. (34) demonstrated dramatic brain transcriptome remodeling in the human brain during postnatal development and these developmental changes are delayed relative to other primates. The delay was not uniform across the human transcriptome but affected a specific subset of genes that play a potential role in neural development. The authors propose that this gene-specific transcriptional neoteny delayed timing of human sexual development which, in turn, may have caused enhanced cognitive function in humans compared to other primates (34). From the current dataset, we cannot distinguish whether the neotenic patterns we observed represent a wholesale shift in expression timing across the brain of brood parasites or whether there are differences in the extent and/or timing of delays across distinct brain regions and molecular pathways. In either case, however, the patterns we report here strongly suggests ontogenetic gene expression shifts in the POA have occurred in the evolution of avian brood parasitism.

Specific subsets of neotenic or downregulated genes may play a role in the appearance of the brood parasitic strategy. We therefore further explored differentially expressed transcripts to identify genes potentially involved in maternal-related neural plasticity. Our examination of differentially expressed transcripts yielded 38 candidate genes associated with structural and functional plasticity. Neuromodulators such as mesotocin, AVT, and galanin have a clear and well-documented role in mammalian parental behaviors (21, 71, 72); however, the role of structural-related genes is less well established. For instance, stathmin is a phosphorylation-regulated tubulin-binding protein plays a critical role in regulating the microtubule cytoskeleton and may be required for axon formation during neurogenesis (73). Stathmin knockout mice have deficient innate and learned fear, which leads to inaccurate threat assessment. This, in turn, results in a profound loss in observed maternal behavior in females as they lack motivation for retrieving pups and are unable to choose a safe location for nest-building (73). Similarly, mesencephalic astrocyte-derived neurotrophic factor (MANF) is a neurotrophic factor that selectively promotes the survival of dopaminergic neurons, which is critical for maternal care as dopamine is one of the most potent prolactin inhibiting factors and therefore dopaminergic neurons are involved in the regulation of parental behaviors (21, 58). It is the case, however, the pattern of expression of several candidate genes in our study does not reflect expression patterns reported in other studies (21, 71–72). However, this may be a consequence of compensatory mRNA expression as described above as well as a reduction in receptor abundance, which may also cause a compensatory upregulation of ligands. While the functional outcomes of these modifications in transcript expression are not clear, these genes do provide excellent candidates for future study on mechanisms of brood parasitism and general evolutionary shifts in parental behavior.

While some candidate genes we identified have been in the spotlight for some time, the bulk of these studies focus on adults. Novel behavioral phenotypes, however, may be related to the ways in which these peptides shape neural circuits and influence social processes during development. Therefore, it is also possible that the higher mRNA expression seen in our study is a component of the neotenic brain as it is the case that experience-dependent brain plasticity is elevated during juvenile critical periods and declines into adulthood (74–76). In songbirds, this has been termed the constitutive plasticity hypothesis which was proposed to address gene patterns during song recognition in songbirds (76). According to this hypothesis, heightened information storage during critical developmental periods is associated with sustained gene expression which may enhance sensitivity to song tutoring, whereas gene expression becomes suppressed in adults and is only induced when the adult bird experiences a salient social stimulus (76). Consequently, it is possible there may be a transferal from “constitutive plasticity” in juveniles to “regulated plasticity” in adults (76), as has been proposed to explain song learning in birds (74). Indeed, in some cases, these critical developmental periods with elevated neural plasticity are mirrored by elevated mRNA expression patterns. For instance, heightened mRNA expression in juveniles compared to adults occurs in structural and neuromodulatory genes including microtubule associated protein 7, synucleins (78), and insulin growth factor I (IGF-1; 79), and corticotropin-releasing hormone (CRH; 80). Moreover, binding to oxytocin and vasopressin receptors is higher in juvenile compared to adult rats in a region-specific manner (81). Thus, modification of neuromodulatory activity and structural plasticity across development provides a potential mechanism for the evolution of novel social behaviors across taxa.

Targeted analysis of candidate genes in other key hypothalamic regions reveals that the differential gene expression within the POA is not specific to this region. Four candidate genes including, mesotocin, AVT, galanin and prostaglandin synthase were examined within four additional hypothalamic brain regions that were pooled as a single sample. Three of the four genes were differentially expressed in these additional hypothalamic regions at the 0.05 level; the exception being prostaglandin synthase (p = 0.059). Moreover, three of the four genes followed the same expression pattern within the POA. The only gene that reversed expression pattern outside the preoptic region is AVT. Within additional hypothalamic regions, AVT exhibited a significant knockdown pattern in the brood parasite rather than the elevated expression pattern exhibited in the POA. Differential candidate gene expression patterns in additional hypothalamic regions indicates that other brain regions may have been the targets of evolution during the transition to brood parasitic strategies, rather than simply the POA. This supports additional studies that reveal hypothalamic regions in female cowbirds exhibit volumetric and neurogenesis differences in female cowbirds in comparison to male cowbirds and red-winged blackbirds (82,83,84). Considerable modifications have also been demonstrated in the auditory physiology of cowbirds (85) Together these studies, along with the results presented here, indicated evolution may have produced wholesale transformations in neural architecture to produce an animal that could successfully navigate the challenges of the brood parasitic strategy.

Despite clear changes in circulating hormone levels following prolactin treatment, POA gene expression in our brood parasitic species was remarkably insensitive to prolactin treatment. We found only four transcripts DE due to prolactin treatment in cowbirds. Previous studies in brown-headed cowbirds have found similar results from a behavioral perspective. That is, prolactin administered to female cowbirds induces younger females to exhibit incubation-like behaviors more often in prolactin treated birds as compared to older females, indicating sensitivity to prolactin changes across development with adult female cowbirds becoming prolactin insensitive (86). This prolactin insensitivity may be related to the lower levels of prolactin receptor expression in the POA in parasitic brown-headed cowbirds as compared to non-parasitic red-winged blackbirds (20). While prolactin receptor abundance in our study was not significantly different with FDR corrections between parasitic and non-parasitic species, prolactin receptor transcript levels are lower in cowbirds, and particularly brown-headed cowbirds, than in red-winged blackbirds (Fig. S1). Thus, prolactin receptor transcript abundance reported here reflects the pattern of prolactin receptor protein reported by Ball, 1991 (20). Future studies will investigate prolactin and non-prolactin treatments in parasitic and non-parasitic birds to determine if prolactin receptor downregulation in the POA serves as the gateway to prolactin transcriptional insensitivity in this brain region.

Our results provide novel contributions to our understanding neurobiological-basis of brood parasitism. We identify a set of CDE genes that may be conserved across brood parasitic species of different lineages and may underlie transition to the brood parasitic lifestyle in multiple lineages of parasites. In addition, we demonstrate largely neotenic patterns of gene expression in these CDE genes. Finally, neotenic expression patterns extend to candidate genes underlying neuromodulation, structural plasticity, and maternal behavior. Together, these results provide a solid foundation for future investigations of neural mechanisms mediating the emergence of brood parasitism and the evolution of novel social behavioral phenotypes across taxonomic groups.

## Acknowledgements

This study was funded by Texas EcoLabs grants provided to KSL. Support in the field was provided by Balcones Wildlife Refuge as well as Drs. Chris Elliot and Jean Deo. We also thank Michael Brewer for developing code for ortholog identification and alignment, Caitlin Friesen for help with animal care, Drs. Mark Hauber, Mary Ramsey and Kim Hoke for helpful discussions of this work.

## Supplemental figures

**Supplemental figure 1.**
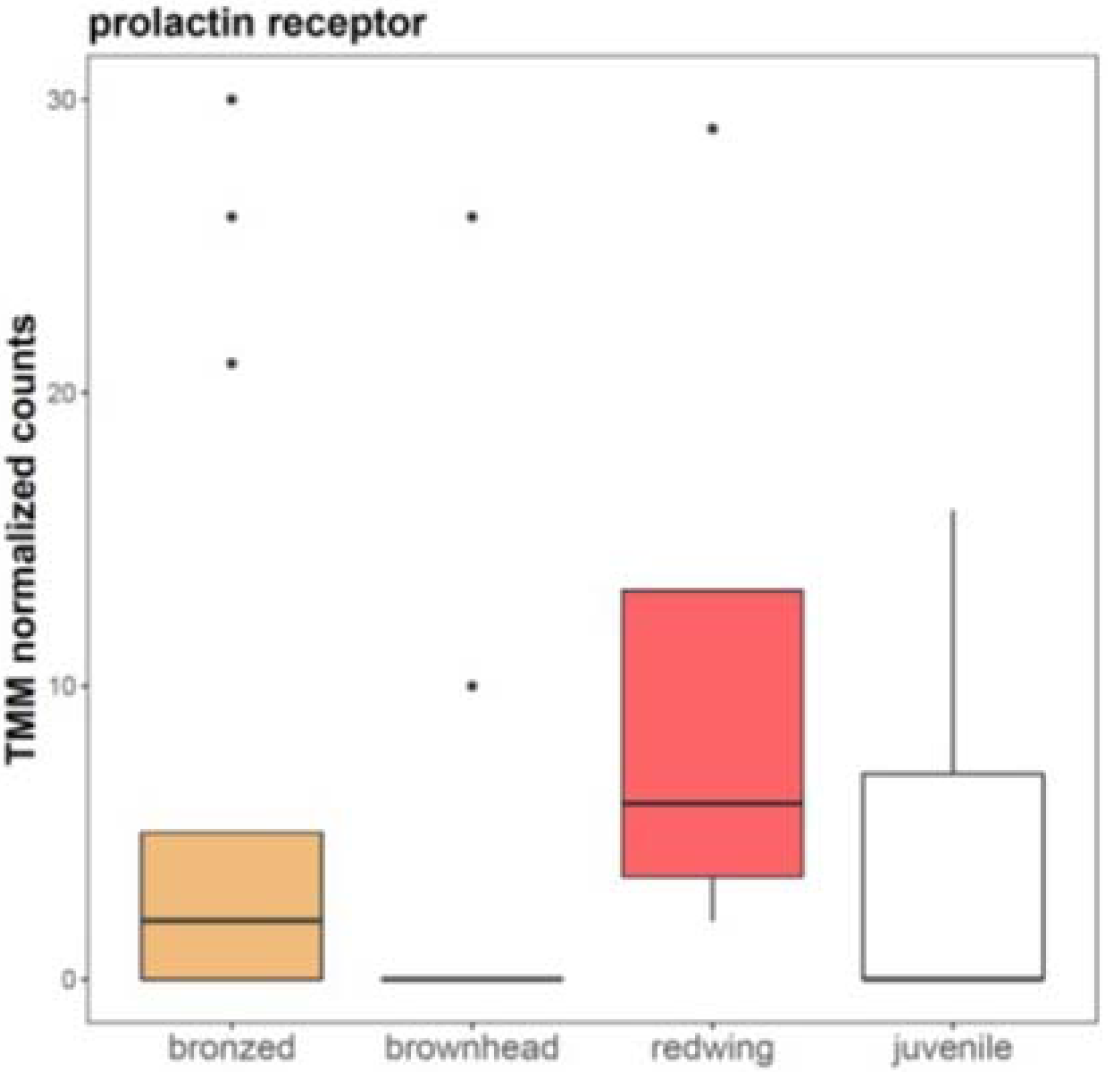
Comparison of prolactin receptors between parasitic cowbirds and non-parasitic adult and juvenile red-winged blackbirds. All error bars represent SEM.

**Supplemental table 2.**
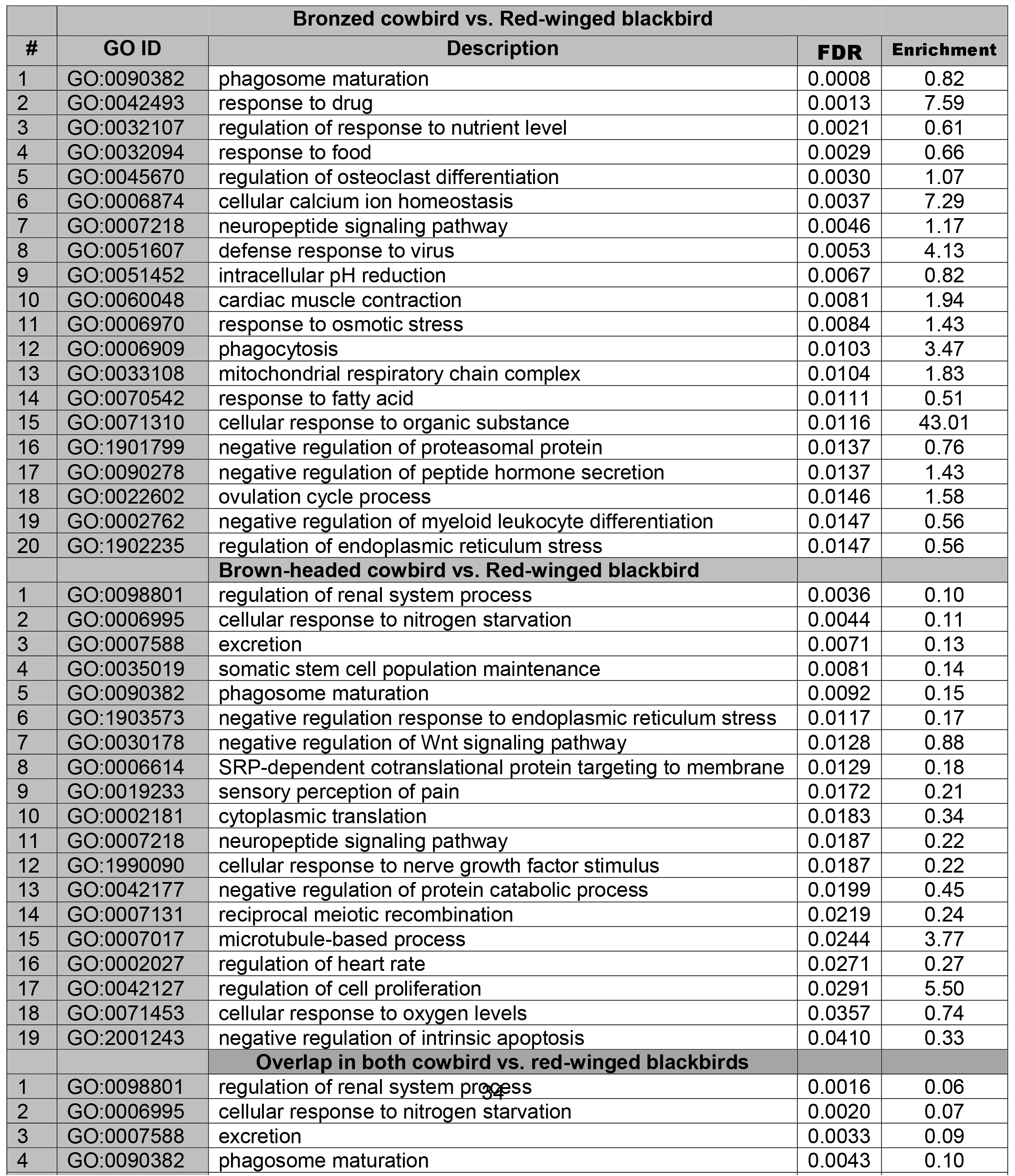
List of differentially expressed genes with a 0.1 cutoff. Comparisons in which DE genes were identified include bronzed cowbird vs red-winged blackbird, brownheaded cowbird vs. red-winged blackbird, brown-headed vs. bronzed cowbirds and genes with concordant differential expression (CDE) between cowbirds when compared to red-winged blackbirds. The pattern of gene regulation is described for CDE genes.

**Supplemental table 3.**
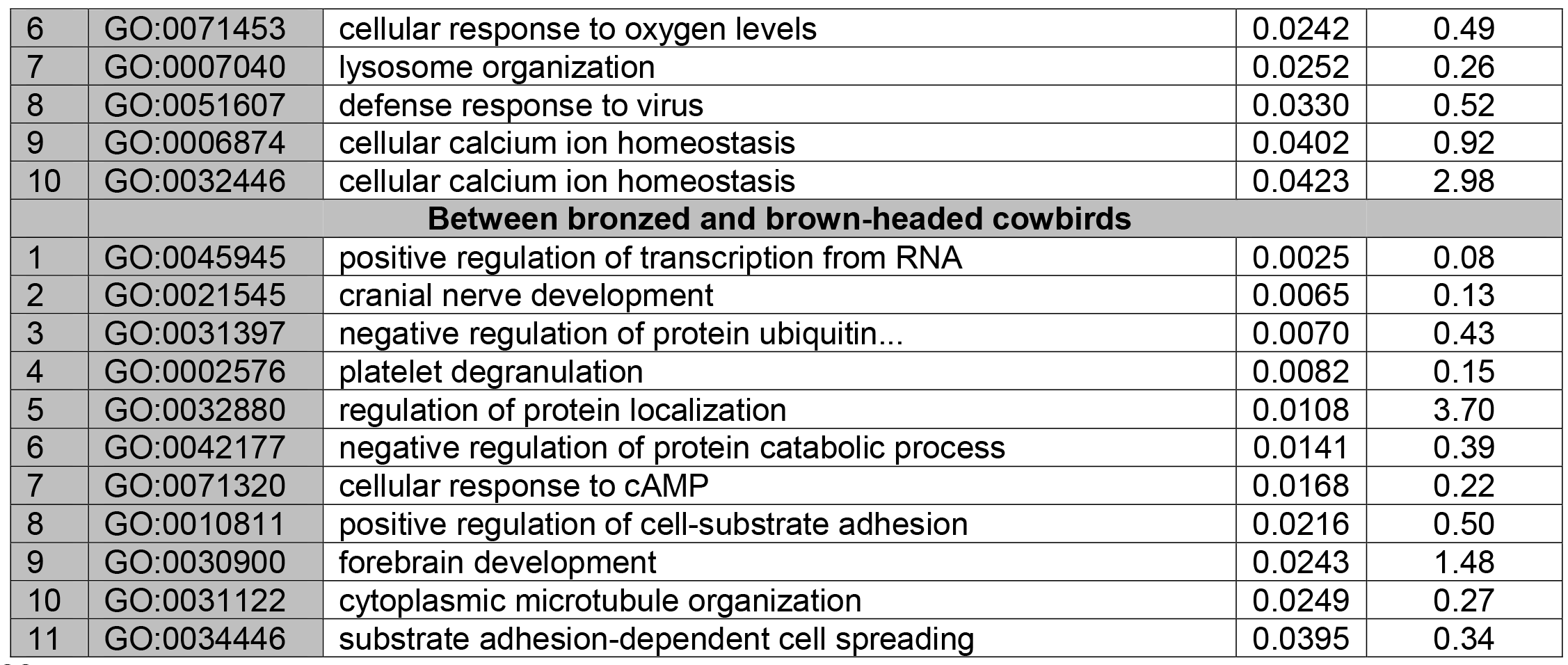
Top 20 gene ontology (GO) categories are listed in four pairwise comparisons: Bronzed cowbirds compared to red-winged blackbirds, brown-headed cowbirds compared to red-winged blackbirds, overlap between both cowbirds and red-winged blackbirds and between brown-headed and bronzed cowbirds.

**Supplemental table 4.** List of differentially expressed genes (DE) in a comparison between adult and juvenile red-winged blackbirds

| # | ID | Red-winged blackbird adult vs. juvenile |  |
| --- | --- | --- | --- |
|  |  | Description | FDR |
| 1 | CART | Cocaine- and amphetamine-regulated transcript protein | 4.14x 10<sup>-6</sup> |
| 2 | GLO2 | Hydroxyacylglutathione hydrolase, mitochondrial | 0.0004 |
| 3 | SELP | Selenoprotein Pb | 0.0019 |
| 4 | NLGX | Neurologin-4, X-linked | 0.00197 |
| 5 | FEZF1 | Fez family zinc finger protein 1 | 0.00197 |
| 6 | IGLL5 | Immunoglobulin lambda-like polypeptide 5 | 0.0027 |
| 7 | XPO7 | Exportin-7 | 0.0027 |
| 8 | NDOR1 | NADPH-dependent diflavin oxidoreductase 1 | 0.0058 |
| 9 | HMX3 | Homeobox protein HMX3 | 0.0060 |
| 10 | HMBX1 | Homeobox-containing protein 1 | 0.0086 |
| 11 | MYPR | Myelin proteolipid protein | 0.0109 |
| 12 | IDS | Iduronate 2-sulfatase | 0.0163 |
| 13 | S10AB | Protein S100-A11 | 0.0319 |
| 14 | EMC6 | ER membrane protein complex subunit 6 | 0.0464 |
| 15 | VIP | VIP peptides | 0.0464 |

**Supplemental table 5.**
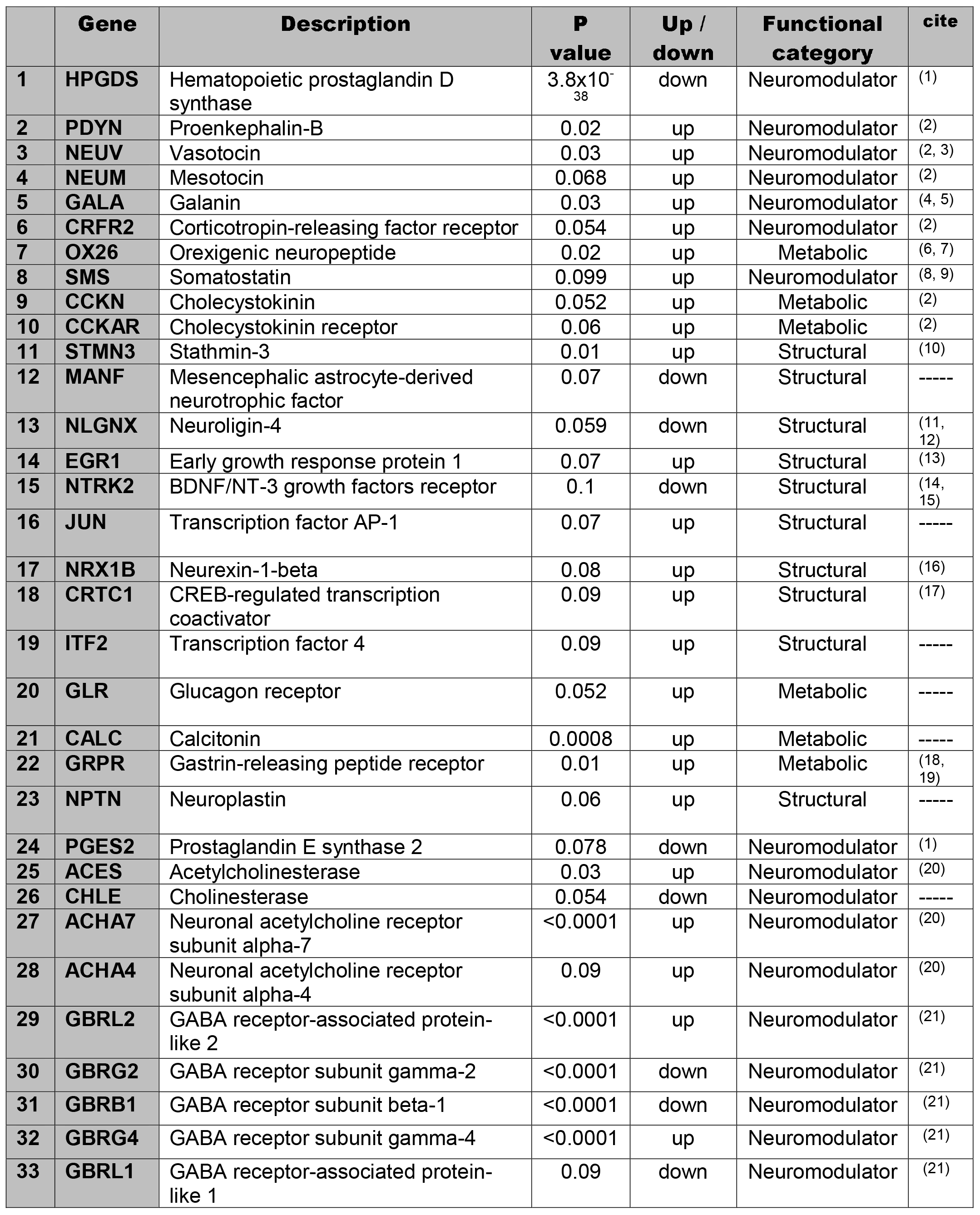
Differentially expressed candidate genes between at least one parasitic species and the non-parasitic species. These genes are identified as being involved in structural and functional plasticity. Functional plasticity includes neuromodulatory genes and metabolic-related regulators. Structural genes are involved in cellular renovations. Up/down refers to the direction of the expression change in the cowbirds. Citations are listed for each gene with an established role in social or maternal behaviors.

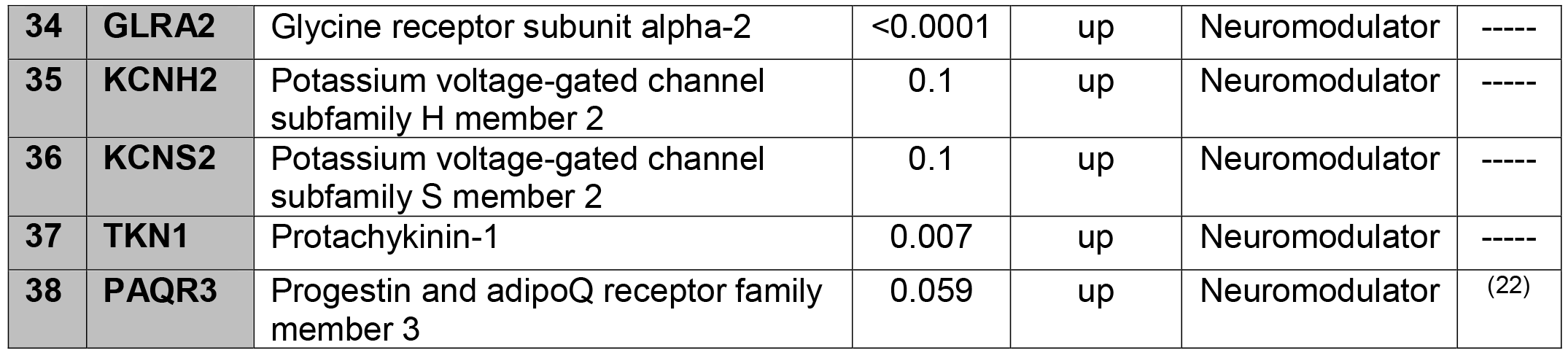

## References

1. Clutton-Brock TH. 1991 The evolution of parental care. Princeton University Press.

2. Klug H, Bonsall MB. 2010 Life history and the evolution of parental care. Evol. 64, 823–835.

3. Canon JB, Vannier J. 2016 Waptia and the diversification of brood care in early Arthropods. Curr. Biol. 26, 69–74.

4. Alonso-Alvarez C, Velando A. 2012 Benefits and costs of parental care. The Evolution of Parental Care. Oxford University Press.

5. Klug H, Bonsall MB., Alonzo SH. 2013 The origin of parental care in relation to male and female life history. Ecol. Evol. 3, 779–791.

6. Lack DL. 1968 Ecological Adaptations for Breeding in Birds. Chapman and Hall.

7. Cockburn A. 1998 Evolution of Helping Behavior in Cooperatively Breeding Birds. Ann. Rev. Ecol. Syst. 29, 141–177.

8. Cockburn A. 2006 Prevalence of different modes of parental care in birds. Proc. R Soc. Lond. B Biol. Sci. 273, 1375–1383.

9. Winfree R. 1999 Cuckoos, cowbirds and the persistence of brood parasitism. Trends Ecol. Evol. 14, 338–343.

10. Lanyon SM. 1992 Interspecific brood parasitism in blackbirds (Icterinae): a phylogenetic perspective. Science 255, 77–79.

11. Davies NB. 2000 Cuckoos, Cowbirds and Other Cheats. T & AD Poyser Ltd, London.

12. Sorenson MD, Payne RB. 2002 Molecular genetic perspectives on avian brood parasitism. Integr. Comp. Biol. 42, 388–400.

13. Sato T. 1986 A brood parasitic catfish of mouthbrooding cichlid fishes in Lake Tanganyika. Nature 323, 58–59.

14. Davies NB, Bourke AFG, Brooke M de L. 1989 Cuckoos and parasitic ants: Interspecific brood parasitism as an evolutionary arms race. Trends. Ecol. Evol. 4, 274–278.

15. Zink AG. 2000 The Evolution of intraspecific brood parasitism in birds and insects. Am. Nat. 155, 395–405.

16. Hamilton WJ, Orians GH. 1965 Evolution of brood parasitism in altricial birds. Condor 67, 361–382.

17. Payne RB. 1977 The ecology of brood parasitism in birds. Ann. Rev. Ecol. Syst. 8, 1–28.

18. Rothstein SI. 1993 An experimental test of the Hamilton-Orians hypothesis for the origin of avian brood parasitism. Condor 95, 1000–1005.

19. Semel B, Sherman PW. 2001 Intraspecific parasitism and nest-site competition in wood ducks. Anim. Behav. 61, 787–803.

20. Ball GF. 1991 Endocrine mechanisms and the evolution of avian parental care. Acta XX Congr. Int. Ornithol., 984–991.

21. Numan M, Insel TR. 2006 The Neurobiology of Parental Behavior, Springer.

22. Adkins-Regan E. 2013 Hormones and Animal Social Behavior, Princeton University Press.

23. Chokchaloemwong D, et al. 2013 Mesotocin and maternal care of chicks in native Thai hens (Gallus domesticus). Horm. Behav. 64, 53–69.

24. Rivas M, Torterolo P, Ferreira A, Benedetto L. 2016 Hypocretinergic system in the medial preoptic area promotes maternal behavior in lactating rats. Peptides 81, 9–14.

25. Dufty AM, Goldsmith AR, Wingfield JC. 1987 Prolactin secretion in a brood parasite, the brownheaded cowbird, Molothrus ater. J. Zool. 212, 669–675.

26. Buntin JD. 1996 Neural and hormonal control of parental behavior in birds. Adv. Study Behav. 25, 161–213.

27. Buntin JD, Buntin L. 2014 Increased STAT5 signaling in the ring dove brain in response to prolactin administration and spontaneous elevations in prolactin during the breeding cycle. Gen. Comp. Endocrinol. 200, 1–9.

28. Smiley KO, Adkins-Regan E. 2016 Prolactin is related to individual differences in parental behavior and reproductive success in a biparental passerine, the zebra finch (Taeniopygia guttata). Gen. Comp. Endocrinol. 234, 88–94.

29. Smiley KO, Adkins-Regan E. 2016 Prolactin is related to individual differences in parental behavior and reproductive success in a biparental passerine, the zebra finch (Taeniopygia guttata). Gen. Comp. Endocrinol. 234, 88–94.

30. Smiley KO, Adkins-Regan E (2018) Loering prolactin reduces post-hatch parental care in male and female zebra finches (Taeniopygia guttata). Horm. Behav. 98, 103–114.

31. Payne RB. 1967 Interspecific communication signals in parasitic birds. Am. Nat. 101, 363–375.

32. Zákány J, Gérard M, Favier B, Duboule D. 1997 Deletion of a HoxD enhancer induces transcriptional heterochrony leading to transposition of the sacrum. EMBO J 16, 4393–4402.

33. Kim J, Kerr JQ, Min GS. 2000 Molecular heterochrony in the early development of Drosophila. Proc. Natl. Acad. Sci. U S A 97, 212–216.

34. Somel M, et al. 2009 Transcriptional neoteny in the human brain. Proc. Natl. Acad. Sci. 106, 5743–5748.

35. Buntin JD, Hnasko RM, Zuzick PH. 1999 Role of the ventromedial hypothalamus in prolactin-induced hyperphagia in ring doves. Physiol. Behav. 66, 255–261.

36. Wang Q, Buntin JD. 1999 The roles of stimuli from young, previous breeding experience, and prolactin in regulating parental behavior in ring doves (Streptopelia risoria). Horm. Behav. 35, 241–253.

37. Lynch KS, Wilczynski W. 2006 Social regulation of plasma estradiol concentration in a female anuran. Horm. Behav. 50, 101–106.

38. Ewing B, Green P. 1998 Base-calling of automated sequencer traces using phred. II. Error probabilities. Genome Res. 8, 186–194.

39. Lunter G, Goodson M. 2011 Stampy: a statistical algorithm for sensitive and fast mapping of Illumina sequence reads. Genome Res. 21, 936–939.

40. Song L, Florea L. 2015 Rcorrector: efficient and accurate error correction for Illumina RNA-seq reads. GigaScience 4, 48.

41. Grabherr MG, et al. 2011 Full-length transcriptome assembly from RNA-Seq data without a reference genome. Nat. Biotechnol. 29, 644–652.

42. Haas BJ, et al. 2013 De novo transcript sequence reconstruction from RNA-seq using the Trinity platform for reference generation and analysis. Nat. Protoc. 8, 1494–1512.

43. Bryant DM, et al. 2017. A Tissue-mapped Axolotl de novo transcriptome enables identification of limb regeneration factors. Cell. Rep. 18, 762–776.

44. Simão FA, Waterhouse RM, Ioannidis P, Kriventseva EV, Zdobnov EM. 2015 BUSCO: assessing genome assembly and annotation completeness with single-copy orthologs. Bioinforma Oxf. Engl. 3, 3210–3212.

45. MacManes MD. 2016 Establishing evidenced-based best practice for the de novo assembly and evaluation of transcriptomes from non-model organisms. bioRxiv:035642.

46. Moreno-Hagelsieb G, Latimer K. 2008 Choosing BLAST options for better detection of orthologs as reciprocal best hits. Bioinforma Oxf. Engl. 24, 319–324.

47. Katoh K, Standley DM. 2013 MAFFT multiple sequence alignment software version 7: improvements in performance and usability. Mol. Biol. Evol. 30, 772–780.

48. Suyama M, Torrents D, Bork P. 2006 PAL2NAL: robust conversion of protein sequence alignments into the corresponding codon alignments. Nucleic Acids Res. 34:W609–612.

49. Kück P. 2009 ALICUT: a Perlscript which cuts ALISCORE identified RSS. Zool Forschungsmuseum Alexander Koenig ZFMK.

50. Bray NL, Pimentel H, Melsted P, Pachter L. 2016 Near-optimal probabilistic RNA-seq quantification. Nat. Biotechnol. 34, 525–527.

51. Love MI, Huber W, Anders S. 2014 Moderated estimation of fold change and dispersion for RNA-seq data with DESeq2. Genome Biol. 15, 550.

52. Alexa A, Rahnenführer J. 2009 Gene set enrichment analysis with topGO Available at: https://bioconductor.riken.jp.

53. Levy F, Gheusi G, Keller M. 2011 Plasticity of the parental brain: a case for neurogenesis. J Neuroendocrinol 23, 984–993.

54. Numan M, Sheehan TP. 1997 Neuroanatomical circuitry for mammalian maternal behavior. Ann. N.Y. Acad. Sci. 807, 101–125.

55. Numan M, Stolzenberg DS. 2009 Medial preoptic area interactions with dopamine neural systems in the control of the onset and maintenance of maternal behavior in rats. Front. Neuroendocrinol. 30, 46–64.

56. Theodosis DT, Pierre K, Poulain DA. 2000 Differential expression of two adhesion molecules of the immunoglobulin superfamily, F3 and polysialylated NCAM, in hypothalamic magnocellular neurones capable of plasticity. Exp. Physiol. 85 Spec No:187S–196S.

57. Shams S, et al. 2012 Dendritic morphology in the striatum and hypothalamus differentially exhibits experience-dependent changes in response to maternal care and early social isolation. Behav. Brain. Res. 233, 79–89.

58. Stolzenberg DS, et al. 2007 Dopamine D1 receptor stimulation of the nucleus accumbens or the medial preoptic area promotes the onset of maternal behavior in pregnancy-terminated rats. Behav. Neurosci. 121, 907–919.

59. Lynch KS, Ramsey ME and Cummings ME. 2012 The mate choice brain: comparing gene profiles between female choice and male coercive poeciliids. Genes, Brain and Behav. 11, 222–229.

60. Aubin-Horth N, Landry CR, Letcher BH and Hofmann HA. 2005 Alternative life histories shape brain gene expression profiles in males of the same population. Proc Roy Soc London B: Biological Sciences. 272, 1655–1662.

61. Cummings ME, Larkins-Ford J, Reilly CR, Wong RY, Ramsey M and Hofmann HA. 2008 Sexual and social stimuli elicit rapid and contrasting genomic responses. ProcRoy Soc London B: Biological Sciences. 275, 393–402.

62. Hashimoto JG, Beadles-Bohling AS and Wiren KM. 2004 Comparison of RiboGreen^®^ and 18S rRNA quantitation for normalizing real-time RT-PCR expression analysis. Biotechniques. 36, 54–60.

63. Demski LS, Knigge KM. 1971 The telencephalon and hypothalamus of the bluegill (Lepomis macrochirus): evoked feeding, aggressive and reproductive behavior with representative frontal sections. J. Comp. Neurol. 143, 1–16.

64. Schoech SJ, Ketterson ED, Nolan V, Sharp PJ, Buntin JD. 1998 The effect of exogenous testosterone on parental behavior, plasma prolactin, and prolactin binding sites in darkeyed juncos. Horm. Behav. 34, 1–10.

65. Trainor BC, Bird IM, Alday NA, Schlinger BA, Marler CA. 2003 Variation in aromatase activity in the medial preoptic area and plasma progesterone is associated with the onset of paternal behavior. Neuroendocrinology 78, 36–44.

66. Ruscio MG, Adkins-Regan E, 2004 Immediate early gene expression associated with induction of brooding behavior in Japanese quail. Horm. Behav. 46, 19–29.

67. Pereira M, Morrell JI. 2011 Functional mapping of the neural circuitry of rat maternal motivation: effects of site-specific transient neural inactivation. J. Neuroendocrinol. 23, 1020–1035.

68. Liu Y, Beyer A, Aebersold R. 2016 On the Dependency of Cellular Protein Levels on mRNA Abundance. Cell 165, 535–550.

69. Shaw P, et al. 2007 Attention-deficit/hyperactivity disorder is characterized by a delay in cortical maturation. Proc. Natl. Acad. Sci. U.S.A 104, 19649–19654.

70. Kur-Piotrowska A, et al. 2017 Neotenic phenomenon in gene expression in the skin of Foxn1-deficient (nude) mice − a projection for regenerative skin wound healing. BMC Genomics 18, 56.

71. Dulac C, O’Connell LA, Wu Z. 2014 Neural control of maternal and paternal behaviors. Science 345, 765–770.

72. Bendesky A, et al. 2017 The genetic basis of parental care evolution in monogamous mice. Nature 544, 434–439.

73. Martel G, Nishi A, Shumyatsky GP. 2008 Stathmin reveals dissociable roles of the basolateral amygdala in parental and social behaviors. Proc. Natl. Acad. Sci. U.S.A 105, 14620–14625.

74. Jin H, Clayton DF. 1997 Localized changes in immediate-early gene regulation during sensory and motor learning in zebra finches. Neuron 19, 1049–1059.

75. Hübener M, Bonhoeffer T. 2014 Neuronal Plasticity: Beyond the Critical Period. Cell 159, 727–737.

76. London SE, Dong S, Replogle K, Clayton DF. 2009 Developmental shifts in gene expression in the auditory forebrain during the sensitive period for song learning. Dev. Neurobiol. 69, 437–450.

77. Garner CC, Matus A. 1988 Different forms of microtubule-associated protein 2 are encoded by separate mRNA transcripts. J. Cell. Biol. 106, 779–783.

78. Malatynska E, et al. 2006 Levels of mRNA coding for alpha-, beta-, and gamma-synuclein in the brains of newborn, juvenile, and adult rats. J. Mol. Neurosci. MN 29, 269–277.

79. Chan CB, Tse MCL, Cheng CHK. 2006 Regulation and Mechanism of Growth Hormone and Insulin-like Growth Factor-I Biosynthesis and Secretion. The Somatotrophic Axis in Brain Function, Academic Press, Burlington.

80. Romeo RD. 2013 The Teenage Brain: The Stress Response and the Adolescent Brain. Curr. Dir. Psychol. Sci. 22, 40–145.

81. Smith CJW, et al. 2017 Age and sex differences in oxytocin and vasopressin V1a receptor binding densities in the rat brain: focus on the social decision-making network. Brain Struct. Funct. 222, 981–1006.

82. Sherry DF, Forbes MR, Khurgel M.and Ivy GO. 1993 Females have a larger hippocampus than males in the brood-parasitic brown-headed cowbird. PNAS. 90, 7839–7843.

83. Guigueno MF, MacDougall-Shackleton SA and Sherry DF. 2016 Sex and seasonal differences in hippocampal volume and neurogenesis in brood-parasitic brownheaded cowbirds (Molothrus ater). Dev neurobio. 76, 1275–1290.

84. Lynch KS. 2018 Region-specific neuron recruitment in the hippocampus of brown-headed cowbirds Molothrus ater (Passeriformes: Icteridae). Eur Zool Jour. 85, 46–54.

85. Lynch KS, Gaglio A, Tyler E, Coculo J, Louder MI and Hauber ME. 2017 A neural basis for password-based species recognition in an avian brood parasite. J Exp Biol. 220, 2345–2353.

87. Robinson GG and Warner DW. 1964 Some effects of prolactin on reproductive behavior in the brown-headed cowbird (Molothrus ater). Auk. 81, 315–325.

## References in table s5

1. Juntti SA, et al. (2016) A Neural Basis for Control of Cichlid Female Reproductive Behavior by Prostaglandin F2α. Curr Biol CB 26(7):943–949.

2. Numan M, Insel TR (2006) The Neurobiology of Parental Behavior (Springer).

3. Bendesky A, et al. (2017) The genetic basis of parental care evolution in monogamous mice. Nature 544(7651):434–439.

4. Dulac C, O’Connell LA, Wu Z (2014) Neural control of maternal and paternal behaviors. Science 345(6198):765–770.

5. Wu Z, Autry AE, Bergan JF, Watabe-Uchida M, Dulac CG (2014) Galanin neurons in the medial preoptic area govern parental behaviour. Nature 509(7500):325–330.

6. Harris GC, Wimmer M, Aston-Jones G (2005) A role for lateral hypothalamic orexin neurons in reward seeking. Nature 437(7058):556–559.

7. Sakurai T, Mieda M (2011) Connectomics of orexin-producing neurons: interface of systems of emotion, energy homeostasis and arousal. Trends Pharmacol Sci 32(8):451–462.

8. Hofmann HA, Fernald RD (2000) Social status controls somatostatin neuron size and growth. J Neurosci OffJ Soc Neurosci 20(12):4740–4744.

9. Trainor BC, Hofmann HA (2006) Somatostatin regulates aggressive behavior in an African cichlid fish. Endocrinology 147(11):5119–5125.

10. Martel G, Nishi A, Shumyatsky GP (2008) Stathmin reveals dissociable roles of the basolateral amygdala in parental and social behaviors. Proc Natl Acad Sci USA 105(38): 14620–14625.

11. Jamain S, et al. (2008) Reduced social interaction and ultrasonic communication in a mouse model of monogenic heritable autism. Proc Natl Acad Sci USA 105(5):1710–1715.

12. Ey E, et al. (2012) Absence of deficits in social behaviors and ultrasonic vocalizations in later generations of mice lacking neuroligin4. Genes Brain Behav 11(8):928–941.

13. Lynch KS, Gaglio A, Tyler E, Coculo J, Louder MI and Hauber ME. (2017) A neural basis for password-based species recognition in an avian brood parasite. J Exp Biol 220(13): 2345–2353.

14. Branchi I, et al. (2006) Early social enrichment shapes social behavior and nerve growth factor and brain-derived neurotrophic factor levels in the adult mouse brain. Biol Psychiatry 60(7):690–696.

15. Berry A, et al. (2015) Decreased Bdnf expression and reduced social behavior in periadolescent rats following prenatal stress. Dev Psychobiol 57(3):365–373.

16. Rabaneda LG, Robles-Lanuza E, Nieto-González JL, Scholl FG (2014) Neurexin dysfunction in adult neurons results in autistic-like behavior in mice. Cell Rep 8(2):338–346.

17. Breuillaud L, et al. (2012) Deletion of CREB-regulated transcription coactivator 1 induces pathological aggression, depression-related behaviors, and neuroplasticity genes dysregulation in mice. Biol Psychiatry 72(7):528–536.

18. Yamada K, Wada E, Wada K (2000) Male mice lacking the gastrin-releasing peptide receptor (GRP-R) display elevated preference for conspecific odors and increased social investigatory behaviors. Brain Res 870(1–2):20–26.

19. Presti-Torres J, et al. (2007) Impairments of social behavior and memory after neonatal gastrin-releasing peptide receptor blockade in rats: Implications for an animal model of neurodevelopmental disorders. Neuropharmacology 52(3):724–732.

20. Karvat G, Kimchi T (2014) Acetylcholine elevation relieves cognitive rigidity and social deficiency in a mouse model of autism. Neuropsychopharmacol Off Publ Am Coll Neuropsychopharmacol 39(4):831–840.

21. Caldji C, Diorio, J, Anisman H and Meaney MJ (2004) Maternal behavior regulates benzodiazepine/GABA A receptor subunit expression in brain regions associated with fear in BALB/c and C57BL/6 mice. Neuropsychopharmacology, 29(7): 1344.

22. Frye CA, Walf AA, Kohtz AS, Zhu Y (2013) Membrane progestin receptors in the midbrain ventral tegmental area are required for progesterone-facilitated lordosis of rats. Horm Behav 64(3):539–545.

